# Vaccine-specific immune responses against *Mycobacterium ulcerans* infection in a low-dose murine challenge model

**DOI:** 10.1101/800250

**Authors:** Kirstie M. Mangas, Andrew H. Buultjens, Jessica L. Porter, Sarah L. Baines, Estelle Marion, Laurent Marsollier, Nicholas J. Tobias, Sacha J. Pidot, Kylie M. Quinn, David J. Price, Katherine Kedzierska, Weiguang Zeng, David C. Jackson, Brendon Y. Chua, Timothy P. Stinear

**Affiliations:** Department of Microbiology and Immunology, Doherty Institute, University of Melbourne; CRCINA, INSERM, Université de Nantes, Université d’Angers, Angers, France; Molekulare Biotechnologie, Fachbereich Biowissenschaften, Goethe-Universität Frankfurt, Frankfurt am Main, Germany; LOEWE Centre for Translational Biodiversity in Genomics (TBG), Germany; Monash Biomedicine Discovery Institute and Department of Biochemistry and Molecular Biology, Monash University, Clayton, VIC 3800, Australia; Victorian Infectious Diseases Reference Laboratory Epidemiology Unit at The Peter Doherty Institute for Infection & Immunity, The University of Melbourne and Royal Melbourne Hospital, VIC 3000, Australia; Centre for Epidemiology & Biostatistics, Melbourne School of Population & Global Health, The University of Melbourne, VIC 3010, Australia

## Abstract

The neglected tropical disease Buruli ulcer (BU) is an infection of subcutaneous tissue with *Mycobacterium ulcerans*. There is no effective BU vaccine. Here, we assessed an experimental prime-boost vaccine in a low-dose murine tail infection model. We used the enoyl-reductase (ER) domain of the *M. ulcerans* mycolactone polyketide synthases electrostatically coupled with a previously described TLR-2 agonist-based lipopeptide adjuvant, R_4_Pam2Cys. Mice were vaccinated and then challenged via tail inoculation with 14-20 colony forming units (CFU) of an engineered bioluminescent strain of *M. ulcerans*. Mice receiving either the experimental ER vaccine or *Mycobacterium bovis* Bacille Calmette-Guérin (BCG) were equally well protected, with both groups faring significantly better than non-vaccinated animals (*p*<0.05). A suite of 29 immune parameters were assessed in the mice at the end of the experimental period. Multivariate statistical approaches were then used to interrogate the immune response data to develop disease-prognostic models. High levels of IL-2 and low IFN**-γ** produced in the spleen best predicted control of infection across all vaccine groups. Univariate logistic regression then revealed vaccine-specific profiles of protection. High titres of ER-specific IgG serum antibodies together with IL-2 and IL-4 in the draining lymph node (DLN) were associated with protection induced by the experimental ER vaccine. In contrast, high titres of IL-6, TNF**-α**, IFN**-γ** and IL-10 in the DLN and low IFN**γ** titres in the spleen were associated with protection following BCG vaccination. This study suggests an effective BU vaccine must induce localized, tissue-specific immune profiles with controlled inflammatory responses at the site of infection.

## Introduction

Buruli ulcer (BU) is a disease primarily of the subcutaneous tissue caused by infection with *Mycobacterium ulcerans*. BU initially presents as redness of the skin that is often accompanied with oedema and swelling. As the disease progresses, oedema may increase or an open ulcer develop (1, 2), the latter is typically characterised by deep undermined edges with a necrotic core comprised of bacteria, dead skin cells and immune cells (3, 4). Ulcers are predominately found on the extremities of the body such as the upper (27% of cases) and lower limbs (70% of cases) (5). The disease is rarely fatal, but if left untreated extensive destruction of subcutaneous tissue can leave victims with significant deformities and lifelong disabilities (3, 6-9).

BU is likely caused when *M. ulcerans* is introduced beneath the skin. This can occur if a region of contaminated skin surface is punctured or by insertion of an object contaminated with the bacteria into subcutaneous tissue (*e.g.* via an insect bite) (10-12). BU-endemic areas include certain regions of West and Sub-Saharan Africa and south eastern Australia (13, 14).

*M. ulcerans* is a slow-growing bacterium, with a doubling time greater than 48 hours (15, 16) making it difficult for early disease diagnosis as symptoms can take between 4-5 months to appear after primary infection (17, 18). If diagnosed early, however, BU can be treated effectively by combination antibiotic therapy (19-21). Unfortunately, in many cases the disease can initially be misdiagnosed as other, more common skin infections. Additionally, a large proportion of BU cases in African countries occur in rural villages and poorer areas with limited or no access to health care, with patients facing disfigurement and permanent disability. Given that diagnoses are delayed and usually occur after a lesion has become relatively advanced and ulceration extensive (22), development of an effective BU vaccine to protect those in highly endemic areas is of paramount importance.

Currently, the only licensed vaccine against mycobacterial infections approved for human use is the *M. bovis*-derived Bacille Calmette-Guérin (BCG) vaccine for prevention of tuberculosis. This vaccine is cross-protective against *M. ulcerans*, but only delays the onset of disease (23-25). Several experimental vaccines have been tested against *M. ulcerans* infection, as summarised in Table 1. Although different animal models have been utilised to study *M. ulcerans* infections including guinea pig, primate, pig and armadillo (10, 23, 26-31), most studies assessing vaccine efficacy have used mice. Vaccines tested in these murine challenge models have included DNA-based, viral-based, protein subunit and whole cell vaccines (25, 32-35) (Table 1). Among the various vaccines, BCG expressing *M. ulcerans* antigens appears to offer the best protective effect against challenge. Hart et al. (36, 37) showed enhanced protection against BU using a recombinant BCG vaccine that expressed *M. ulcerans* Ag85A or recombinant Ag85B-EsxH fusion protein in a mouse footpad challenge model. Whilst improving the immunogenicity of BCG may be a promising route, there are also some drawbacks; exposure to environmental mycobacteria is believed to decrease the efficacy of the BCG vaccine and administration in areas where people have been BCG-exposed may be problematic (38).

**Table 1.** Summary of putative *M. ulcerans* vaccines tested in murine model of BU infection.

| Vaccine type | Description of components | *M. ulcerans* challenge dose | Test for mycolactone production<sup>a</sup> | Challenge model | Efficacy compared to BCG | Ref |
| --- | --- | --- | --- | --- | --- | --- |
| DNA-based | pCDNA3 vector encoding Hsp65 | 10<sup>4</sup> AFB (strain 1615 ATCC35840) | Not mentioned | Tail | Less protective | (52) |
| DNA-based | V1Jns.tPA vector encoding Ag85A | 3 x 10<sup>4</sup> AFB (strain 5150) or 10<sup>5</sup> AFB (strain 04-855) | Not mentioned | Footpad | Less protective | (32, 33) |
| DNA-based | Primary vaccination with V1Jns.tPA plasmid encoding mycolactone polyketide domains and boosted with recombinant domain proteins emulsified in Gerbu adjuvant. | 10<sup>5</sup> AFB (strain 1615) | Not mentioned | Footpad | Less protective | (49) |
| Viral | Vesicular stomatitis virus (VSV) replicon particles expressing *M. ulcerans* codon optimised antigens MUL_2232 and MUL_3720 | 30µl of 2.8 x 10<sup>5</sup> CFU/ml stock (8.4 x 10<sup>3</sup> CFU/dose) (strain S1013) | Not mentioned | Footpad | Less protective | (113) |
| Subunit | MUL2232 and MUL3720 adjuvanted with GLA-SE (EM408) | 1.5 x 10<sup>6</sup> or 1.5 x 10<sup>5</sup> CFU (strain S1013) | Not mentioned | Footpad | Less protective | (34) |
| Live Cell | *M. ulcerans* | 10<sup>6.3</sup> or 10<sup>4.3</sup> viable bacteria | Not mentioned | Footpad | Less protective | (114) |
| Live cell | *M. marinum* | 10<sup>5</sup> bacteria (strain 1615) | Not mentioned | Footpad | More protective<sup>b</sup> | (35) |
| Live cell recombinant | *M. marinum* expressing Ag85A (on vector) | 10<sup>5</sup> bacteria (strain 1615) | Not mentioned | Footpad | More protective<sup>b</sup> | (35) |
| Live cell recombinant | *M. bovis* BCG expressing Ag85A (on vector pMV261) | 10<sup>5</sup> bacteria (strain 1615) | Not mentioned | Footpad | More protective | (36) |
| Live cell recombinant | *M. bovis* BCG expressing Ag85B-EsxH fusion protein Ag85A (on vector pMV261) | 10<sup>5</sup> bacteria (strain 1615) | Not mentioned | Footpad | More protective | (37) |
| Inactivated whole cell | Mycolactone-negative *M. ulcerans* (strain 5114) | 4 log<sub>10</sub> or 3 log<sub>10</sub> CFU (strain 98-912) | Not mentioned | Footpad | Less protective | (25) |
| Inactivated whole cell | Mycolactone-deficient attenuated *M. ulcerans* (strain ATCC19423) | 10<sup>6</sup> bacteria (strain TMC1615) | Not mentioned | Footpad | Not compared | (54) |
| Inactivated whole cell | Formalin-treated *M. ulcerans* (strain TMC1615) | 10<sup>6</sup> bacteria (strain TMC1615) | Not mentioned | Footpad | Not compared | (54) |
| Inactivated whole cell | Dewaxed *M. ulcerans* (strain TMC1615) | 10<sup>6</sup> bacteria (strain TMC1615) | Not mentioned | Footpad | Not compared | (54) |
| Phage | Mycobacteriophage D29 (therapeutic vaccine) | 5.5 log<sub>10</sub> AFB (strain 1615) | Not mentioned | Footpad | Not comparable | (53) |

<sup>a</sup> Identifying whether the bacterial culture used for challenge was assessed for mycolactone production before infection.
<sup>b</sup> Vaccine was more protective than the BCG vaccine, however all mice eventually developed footpad swelling.

*M. ulcerans* causes disease primarily through the production of a lipid toxin called mycolactone (39). Mycolactone modulates cell function, in particular secretion of critical cytokines by specifically inhibiting the Sec61 translocon, enabling *M. ulcerans* to escape host immune defences (40-46). The toxin is formed from simple acetate and propionate precursors by three polyketide synthases (PKS) encoded by genes on the plasmid pMUM001 (47, 48). Within each PKS are enzymatic domains that form the mycolactone molecule. Some of these domains have been found to be immunogenic and in particular immune responses against the enoyl reductase (ER) domain have previously been shown to significantly reduce *M. ulcerans* bacterial load in the footpad of a murine prime-boost vaccination study (49). Based on the immunogenic and protective qualities of ER, we have utilised it as a target antigen for a BU subunit vaccine.

Protein antigens generally require an adjuvant to boost immunogenicity and shape immune responses. A known TLR-2 ligand, R_4_Pam_2_Cys, has been previously shown to induce robust antibody responses as well as augmented CD4^+^ and CD8^+^ T cell responses, possibly through promoting dendritic cell antigen uptake and trafficking to lymph nodes (50, 51). Given BU is a disease where the bacteria can be both extracellular and intracellular, the ability of R_4_Pam_2_Cys to robustly engage multiple arms of the adaptive immune system may be beneficial for a BU subunit vaccine.

Previous research has shown that as few as three colony forming units (CFU) of bacteria are required to initiate infection (10), however most animal models challenge with >10^4^ CFU (see Table 1) (25, 32-37, 49, 52-54). This dose is likely to be far higher than the dose of bacteria that leads to natural infection in humans and other animals (10). Such an unrealistic high dose may overwhelm immune responses and underestimate the true efficacy of potential *M. ulcerans* vaccines. For these reasons, we have used a low-dose of *M. ulcerans* in a tail infection challenge model to evaluate an experimental prime-boost subunit vaccine against BU. The experimental subunit vaccine developed here comprised of the *M. ulcerans* mycolactone ER domain protein formulated with the adjuvant, R_4_Pam_2_Cys. Our murine challenge model with physiologically relevant dosing enabled us to more accurately measure vaccine-induced protection and to explore immune correlates of protection against BU.

## Results

### Formulation of the ER-R_4_Pam_2_Cys subunit vaccine candidate

Mycolactone is the key virulence factor produced by *M. ulcerans* and an attractive vaccine target, but the molecule is poorly immunogenic (55). However, the PKS enzymes used by the bacterium to synthesize mycolactones – are highly conserved and immunogenic (49, 56-58). Therefore, we hypothesized that targeting the conserved enzymatic domains of the mycolactone PKS could be an effective vaccine strategy. One domain in particular, the enoyl reductase (ER) protein domain, elicits serum antibodies in BU patients and healthy controls in BU endemic regions (57). The ER protein expressed as an antigen in a DNA-protein prime-boost vaccine has also been shown to reduce bacterial burden in a mouse footpad *M. ulcerans* challenge model (49). Here, we utilised the ER protein to create a novel BU vaccine candidate by electrostatically associating it with the TLR-2 agonist-based lipopeptide R_4_Pam_2_Cys. To formulate this vaccine, recombinant 6xHis-tagged ER protein (37 kDa) was first produced and confirmed by SDS-PAGE (Fig. 1A) and western blotting (Fig. 1B). This protein antigen was formulated with R_4_Pam_2_Cys at various ratios to firstly optimise the formation of protein-lipopeptide complexes. A ratio of 1:1 was sufficient to produce complexes that were larger in size than each constituent on its own (Fig. 1C). While the majority of these complexes were ∼300nm in radius (peak 5), the presence of smaller sized particles of ∼100nm (peak 4) suggests that not all antigen was incorporated within a complex as this size range corresponds to the size of ER protein (peak 3) or R_4_Pam_2_Cys alone (peak 2). Although a 1:3 ratio appears to be more effective for the association of these constituents, the size distribution of form complexes was not monodispersed and appeared as two distinct populations (peak 6 and 7). A 1:5 ratio, however, produced a uniform population of complexes that were ∼500 nm in radius (peak 8) and this formulation was subsequently used to vaccinate animals.

**Figure 1.**
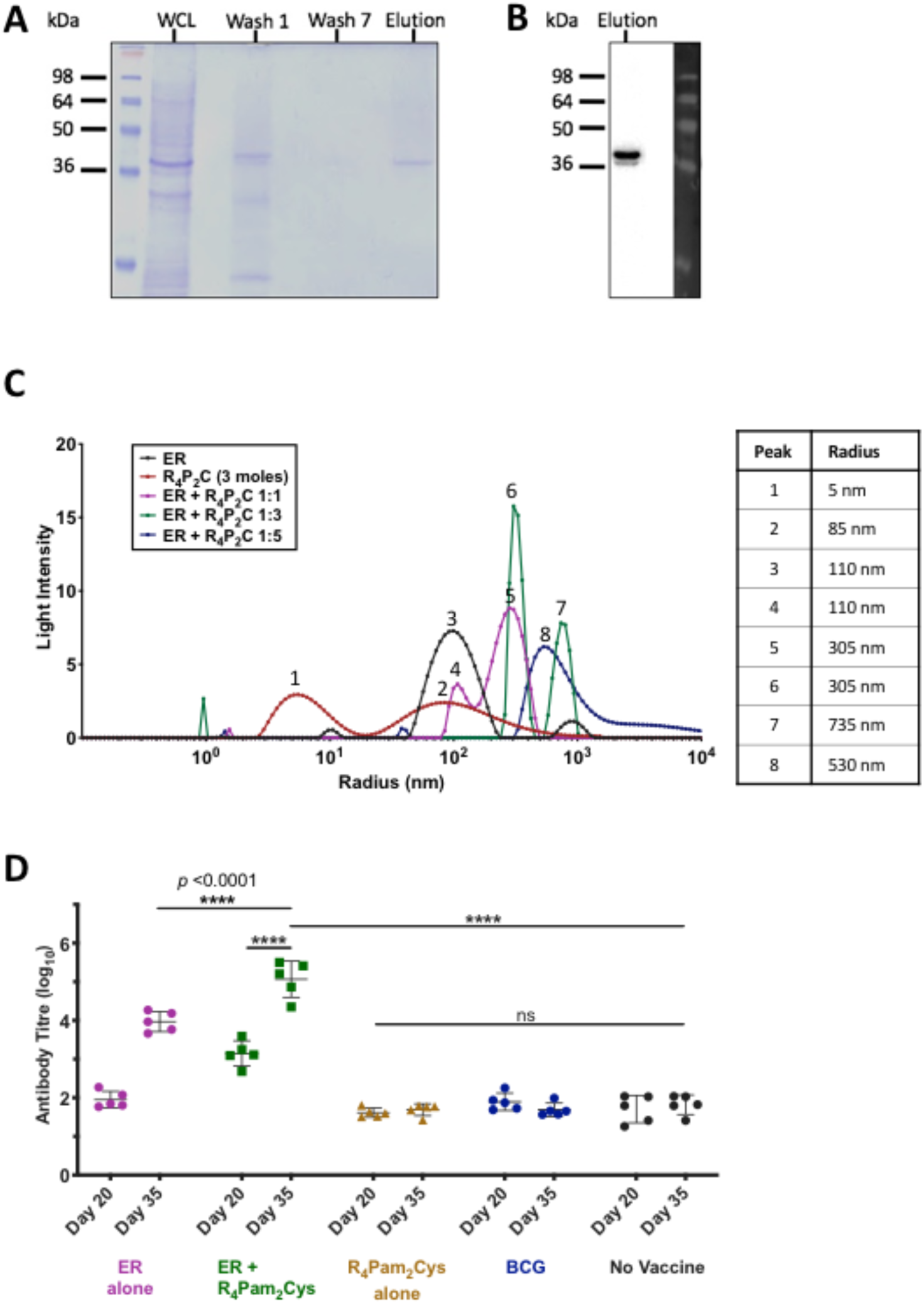
Analysis of purified recombinant ER protein antigen characteristics and formulation with R_4_Pam_2_Cys. **(A)** The presence of recombinant ER protein (∼37 kDa) was monitored by SDS-PAGE at each stage of the purification process; Lane: 1 - Whole cell lysate (WCL), Lane 2 - Wash 1, Lane 3 - Wash 7 and Lane 4 – ER protein elution (containing 10 µg protein). (**B)** Western Blot using an anti-6xHIS-tag antibody to detect the presence of a single band corresponding to the correct molecular weight of the ER protein in the final eluate. **(C)** To analyse the formation of antigen-lipopeptide complexes, a constant amount of antigen was mixed with lipopeptide at different protein:lipopeptide molecular ratios in 50µl of PBS. The size distribution of particles was then analysed by DLS with each profile depicting the hydrodynamic radius (nm) of complexes in each solution. The average radius of each formulation is highlighted in the accompanying table. **(D)** BALB/c mice (n=5/group) were vaccinated on day 0 and day 21 with R_4_Pam_2_Cys alone, ER antigen alone or antigen formulated with R_4_Pam_2_Cys or vaccinated with BCG on day 0 only. Total serum (IgG) antibody against recombinant ER protein were measured by ELISA after the primary dose (day 20) and two weeks after the secondary dose (day 35). Statistical tests were conducted at the 5% significance level. The null hypothesis was rejected if there was a significant difference in mean antibody responses between treatment groups. Note: \**p* < 0.05, \*\**p* < 0.01, \*\*\**p* < 0.001 or \*\*\*\**p* < 0.0001. The error bars represent standard deviation (n=5).

### Evaluation of ER-specific antibody responses in vaccinated mice

To evaluate the ability of the vaccine to induce ER-specific antibody responses, mice were vaccinated, and sera obtained after priming and boosting with the subunit vaccine. Our results showed that primary vaccination with ER + R_4_Pam_2_Cys resulted in significantly higher levels of ER-specific antibody compared to vaccination with unadjuvanted ER antigen (*p*<0.0001) (Fig. 1D). In fact, there was no significant difference in responses between unvaccinated mice and those vaccinated with a single dose of ER alone, R_4_Pam_2_Cys only or BCG. Although a second dose of ER alone was able to increase these responses, the titres were still ∼100-fold less than levels achieved by boosting mice with ER + R_4_Pam_2_Cys (*p*<0.0001). These results not only indicate that ER-specific antibodies can be generated in mice and that the use of R_4_Pam_2_Cys can significantly enhance these responses, but that BCG does not induce cross-reactive antibodies to the ER protein.

### Characterisation of a low-dose M. ulcerans murine tail infection model

We have previously described the use of a low-dose tail infection model for studying insect-mediated transmission of *M. ulcerans* (10). We reasoned that because BU patients were likely to be initially infected with a low bacterial inoculum (10, 11, 59) we could use this model to test the protective efficacy of responses induced by the ER + R_4_Pam_2_Cys vaccine. This model features the use of a bioluminescent strain of *M. ulcerans* (10, 60) and its infectious characteristics are summarised in Fig. 2. Compared to an uninfected tail (Fig 2A and C), sub-cutaneous infection of a tail results in the appearance of a visible ulcer (Fig 2B) exhibiting the highest levels of bioluminescence concentrated around the centre of the lesion (Fig 2D), i.e. where swelling appears to be the greatest, and beginning to diminish around the periphery reflecting a positive correlation between bacterial burden and light emission (61). Histological cross-sections revealed that while tissue integrity of an uninfected tail appears defined and intact (Fig. 2E & F), dramatic differences are observed in the infected tail, typified by loss of muscle, vasculature and epidermis structure and disruptions to surrounding connective tissue (Fig. 2G & H). Further examination of this tissue showed the presence of acid-fast bacilli as well as an infiltration of polymorphonuclear cells (PMNs) (Fig. 2I). Despite evidence of bacteria engulfment by these cells (Fig. 2J), it would appear that this response was insufficient for controlling the infection and preventing disease progression.

**Figure 2.**
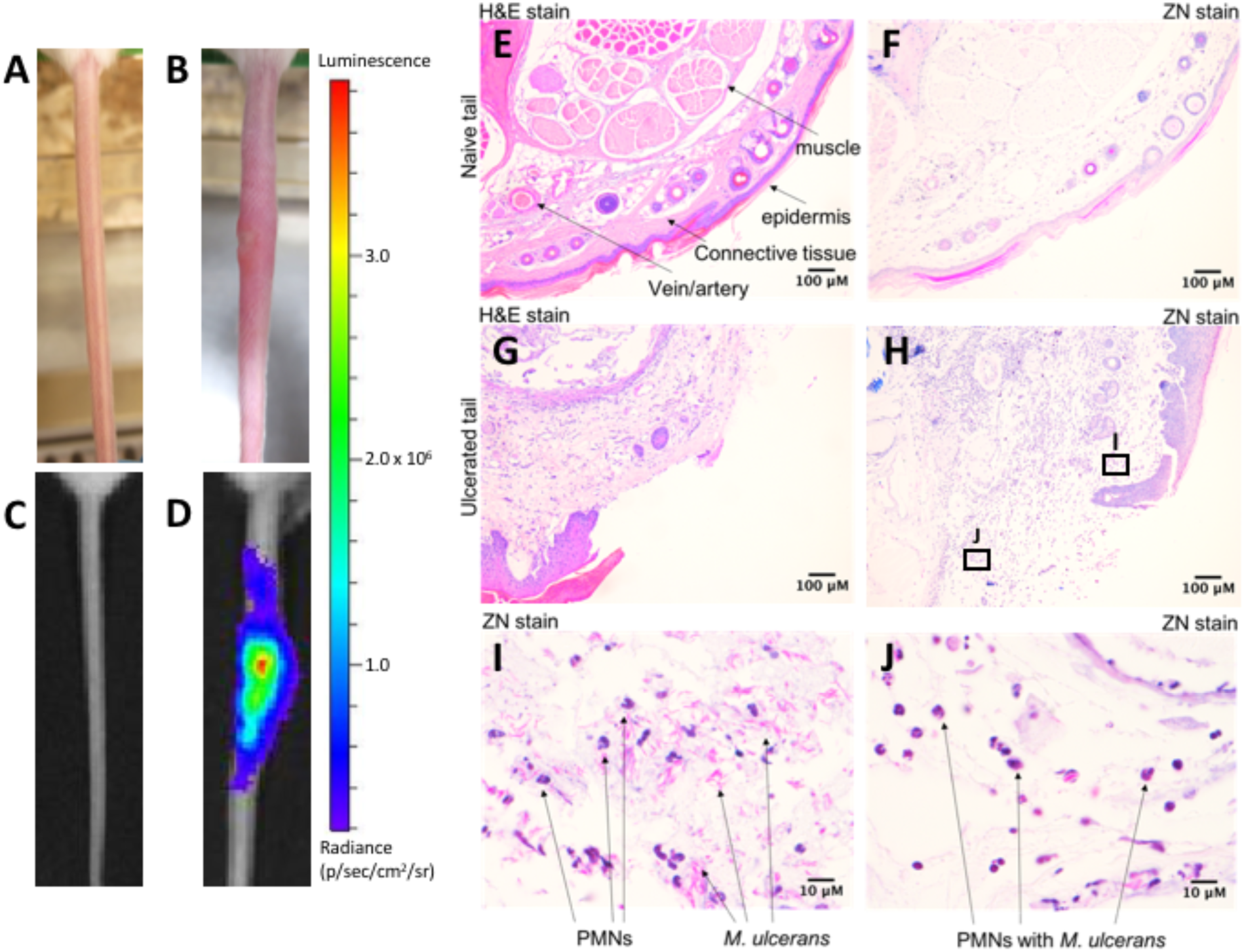
Characterisation of infection using a low-dose bioluminescent *M. ulcerans* strain. Representative light camera images of tails from **(A)** an uninfected BALB/c mouse or **(B)** at the point of ulceration (16 weeks) following intradermal inoculation with 20 CFU of bioluminescent *M. ulcerans*. **(C, D)** The same tails were visualised under an IVIS camera to detect and quantify bioluminescence intensity (as photons/sec). Histological cross section of an **(E, F)** uninfected or **(G, H)** infected tail tissue following haematoxylin & eosin (H&E) and Ziehl-Neelson (ZN) staining. Zoomed images of the regions indicated within the denoted boxes of **(H)** and depicts the presence of polymorphonuclear cells (PMNs) and acid-fast bacilli (ZN staining) within tissue **(I, J)**.

### Monitoring vaccine efficacy using bioluminescent M. ulcerans and in vivo imaging system (IVIS)

We have previously established a correlation between *M. ulcerans* bioluminescence and bacterial burden (61). Here, we set a threshold of ≥ 5×10^5^ photons/second (equivalent to 2.8×10^5^ CFU) as a measure of ineffective disease control, *i.e.* appearance of disease. Mice were vaccinated subcutaneously and then challenged with an estimated 17 CFU (range 14-20 CFU) of bioluminescent *M. ulcerans* via intradermal tail inoculation. Animals were monitored weekly for changes in bioluminescence using IVIS for up to 24 weeks. Fig. 3A shows an example of the progression of bioluminescence (and therefore disease) in an unvaccinated mouse, up to week 16 whereupon the clinical endpoint of the experiment was reached. Bioluminescence for all mice was recorded across the experimental period. Plots for the different treatment groups show the progression in bioluminescence over time (Fig3B-F). Mice from the ER alone, R_4_Pam_2_Cys alone and unvaccinated treatment groups displayed the first detectable bioluminescence at week 7. There also appeared to be threshold in bioluminescence, whereby animals expressing ≥5×10^5^ photons/second from tail lesions became less able to control the infection and progressed to the clinical endpoint (Fig. 3B). The immune response data for all mice is provided (Supplementary Table S1). Using these data, failure-to-protect was defined as tail bioluminescence equal to or greater than 5×10^5^ photons/second at or before week 24 (end of experiment). Therefore, mice were defined as ‘protected’ if bioluminescence was less than 5×10^5^ photons/second at week 24.

**Figure 3.**
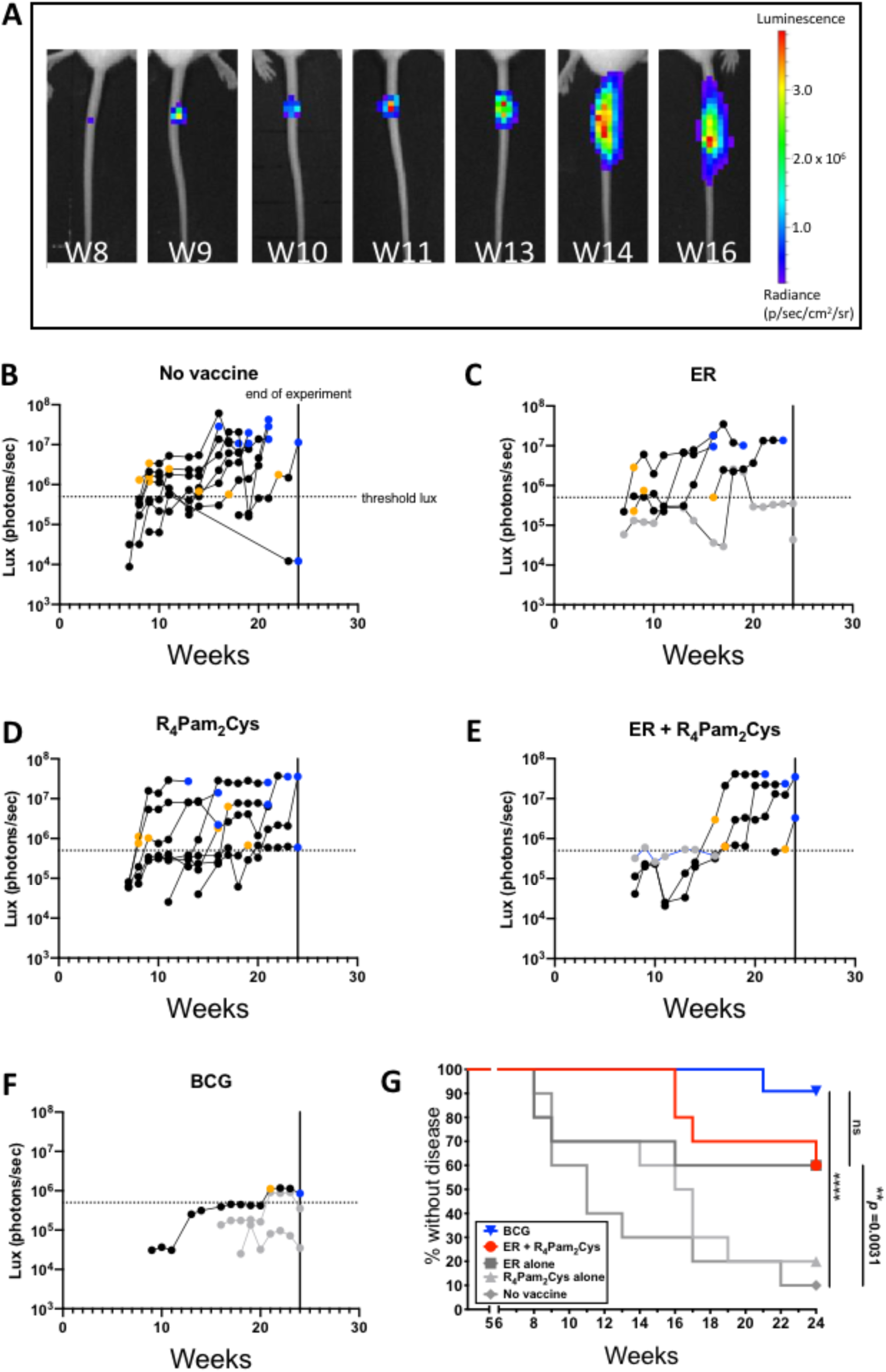
Development of BU over time after vaccination. (A) Tails of mice were intradermally infected with 20 CFU of bioluminescent *M. ulcerans* and imaged weekly by IVIS. Representative panels depict the weekly progression of bioluminescent *M. ulcerans* burden in the tail of an unvaccinated mouse over the course of 16 weeks expressed as photons/second. BALB/c mice (n=10/group) were left **(B)** unvaccinated, or vaccinated on day 0 and day 21 with **(C)** ER antigen alone, **(D)** R_4_Pam_2_Cys alone,, **(E)** ER antigen formulated with R_4_Pam_2_Cys or **(F)** on day 0 with BCG followed by challenge on week 5 with bioluminescent *M. ulcerans*. Threshold bioluminescence (threshold lux) for disease was defined as ≥5×10^5^ photons/second (p/s) as mice that reached this level typically progressed to the clinical (ethical) end point. Mice were classified as diseased if they reached this end point within the 24-weeks following challenge or if their bioluminescence value at week 24 was ≥5×10^5^ p/s. Mice were classified as protected if they did not reach this clinical end point and their bioluminescence value was <5×10^5^ photons/second. The data point depicting when an infected mouse first exhibited bioluminescence at ≥5×10^5^ p/s is represented with a yellow symbol. The data point denoting when a mouse reached clinical endpoint is represented with a blue symbol. Protected mice with detectable bioluminescence are depicted as grey symbols. **G.** Time to bioluminescence measured by IVIS. A survival curve was utilised to analyse the time (weeks) taken for each BU diseased mouse to first reach threshold bioluminescence ≥5×10^5^ photons/second. BCG group (upside down triangle) is labelled in blue, ER + R_4_Pam_2_Cys (circle) is red, and ER alone (square), R_4_Pam_2_Cys alone (upright triangle) and no vaccine (diamond) groups are depicted in grey. Statistical tests were conducted at the 5% significance level. The null hypothesis was rejected if there was a significant difference in survival between groups. Note: \**p* < 0.05, \*\**p* < 0.01, \*\*\**p* < 0.001 or \*\*\*\**p* < 0.0001.

### Vaccination with ER + R_4_Pam_2_Cys offers similar protection to M. bovis BCG vaccine

Survival analysis was conducted to assess vaccine efficacy by measuring the time from infection until tail bioluminescence at the threshold of 5×10^5^ photons/second was reached. Mice that reached this threshold were defined as ‘not protected’. Significantly less ER + R_4_Pam_2_Cys vaccinated mice (4/10 animals) developed disease compared to unvaccinated mice (9/10 animals), indicating ER + R_4_Pam_2_Cys provided some level of protection against disease progression compared to no vaccination (Fig. 3B, E & G) (*p* < 0.01). Mice vaccinated with BCG were best protected with only 1 animal exceeding the bioluminescence threshold (Fig. 3F & G). Although this number of mice was reduced compared to the ER + R_4_Pam_2_Cys vaccinated animals, the difference was not significant (Fig. 3G). However, the bacterial burden (as indicated by mean photon counts/sec at the clinical endpoint) in BCG vaccinated mice was lower than animals that received the ER + R_4_Pam_2_Cys vaccine (means: 6×10^5^ [n=2, range 3.5 – 8.6×10^5^] versus 3.3×10^7^ [n=3, range 2.4 - 4.1×10^7^] photons/sec respectively) suggesting the protective superiority of BCG. There was also no significant difference between the protective efficacy of vaccination with ER alone compared with ER + R_4_Pam_2_Cys, although mice vaccinated with the latter exhibited delayed disease progression (onset at weeks 16-24) compared to ER vaccinated mice (onset at weeks 8-16) indicating that formulation of the antigen with the R_4_Pam_2_Cys adjuvant improved immunity (Fig. 3D, E).

### Measuring immune parameters following vaccination and challenge

At the experimental end-point, sera, spleens and draining lymph nodes (DLN) from all animals were collected and several parameters were further analysed; ER-specific antibodies, CD4^+^ and CD8^+^ T cells and a panel of 11 murine Th1, Th2 and Th17 cytokines. After *M. ulcerans* challenge, mice vaccinated with ER alone or ER + R_4_Pam_2_Cys were found to exhibit significantly more ER-specific antibodies than the other treatment groups (Fig 4A) despite not being fully protected. This indicates that, even though the ER protein is highly immunogenic, anti-ER antibodies do not appear to play a major role in controlling infection.

**Figure 4.**
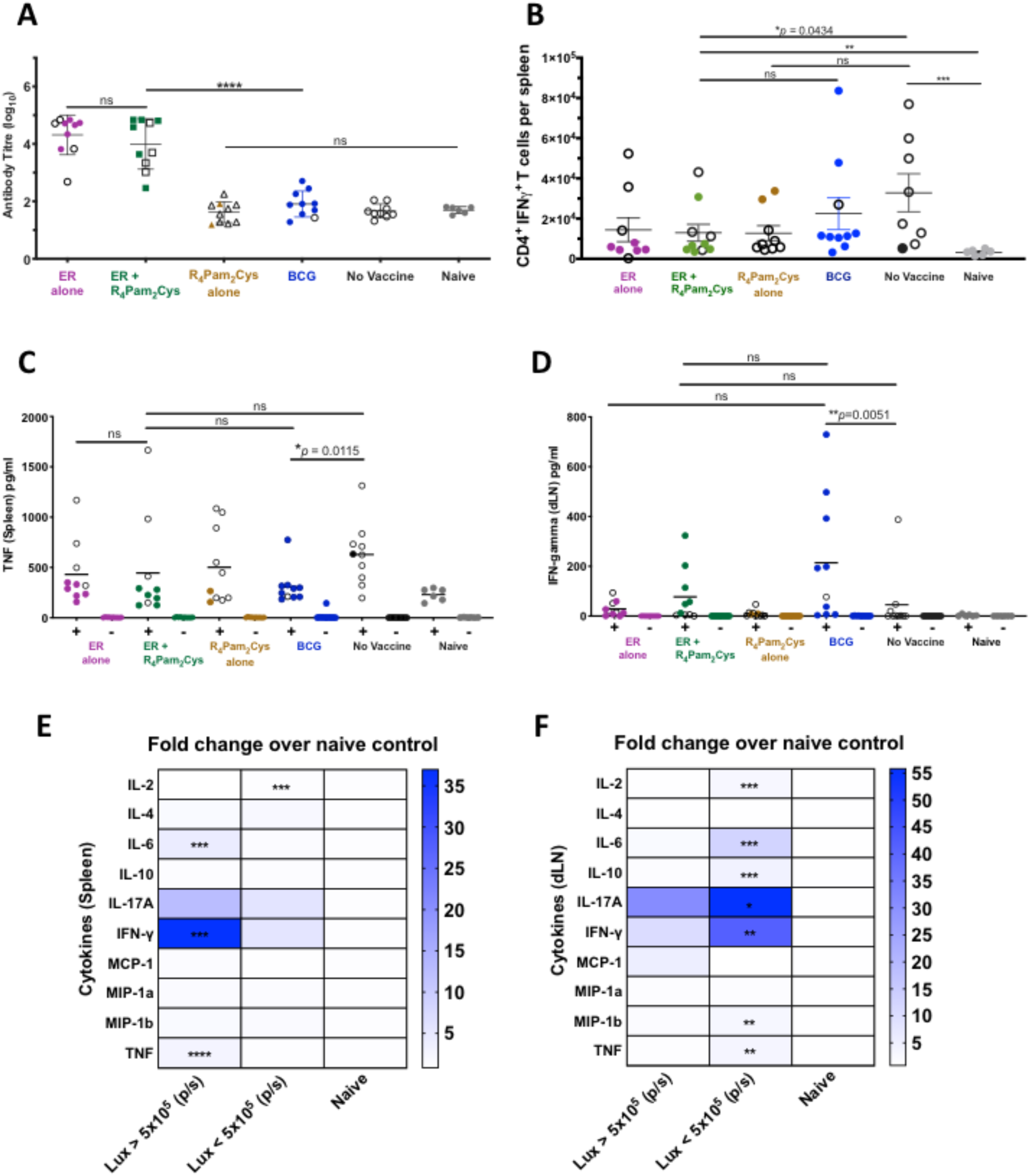
Immune responses after *M. ulcerans* infection. BALB/c mice (n=10/group) were left unvaccinated or vaccinated on day 0 and day 21 with ER antigen alone, ER antigen formulated with R_4_Pam_2_Cys, R_4_Pam_2_Cys alone, or on day 0 with BCG followed by challenge on day 36 with bioluminescent *M. ulcerans*. **(A)** Total serum (IgG) antibody against recombinant ER protein were measured by ELISA after the experimental end point. All data points for diseased mice (bioluminescence ≥5×10^5^ p/s) are depicted with white symbols. Statistical tests were conducted at the 5% significance level. The null hypothesis was rejected if there was a significant difference in mean antibody responses between treatment groups. The error bars represent standard deviation. **(B)** After experimental end point was reached, CD4+ IFN-γ+ T cells were enumerated from the spleen of mice in response to ER protein. The null hypothesis was rejected if there was a significant difference in mean CD4+ IFN-γ+ T cells between treatment groups. Once experimental end point was reached, cytokines from draining lymph nodes and spleens of *M. ulcerans* challenged mice were also measured in response to *in vitro* cell stimulation with recombinant ER protein (Supplementary Table S1). Shown here are cytokine titres **(C)** IFN-γ produced from immune cells in the draining lymph nodes and **(D)** TNF produced from immune cells in the spleen. The null hypothesis was rejected if there was no difference in mean cytokine titres between treatment groups. The black bars represent the mean. Fold change of mean cytokine titres from protected mice (bioluminescence <5×10^5^ p/s) and diseased mice (bioluminescence ≥5×10^5^ p/s) over naïve mice were compared in the (**E)** spleen and (**F)** draining lymph nodes. The null hypothesis was rejected if there was a significant difference in mean cytokine titres between treatment groups. All statistical tests were conducted at the 5% significance level. Note: \**p* < 0.05, \*\**p* < 0.01, \*\*\**p* < 0.001 or \*\*\*\**p* < 0.0001.

We next investigated if there were any differences in the ability of T cells harvested from the spleens of vaccinated mice to produce cytokines following stimulation with recombinant ER protein (Supplementary Table S1). Our results showed that the numbers of IFN-γ producing CD4^+^ T cells across all vaccine groups did not differ, and in some cases, were higher in unvaccinated mice compared to those vaccinated with ER + R_4_Pam_2_Cys vaccine group (Fig 4B) indicating that there was no clear correlation between the frequencies of these cells and protection. Similarly, there also did not appear to be any correlation between TNF-α^+^ CD4^+^ T cells, IFN-γ^+^ CD8^+^ T cells or TNF-α^+^ CD8^+^ T cells and protection (Supplementary Table S1).

Comparing levels of cytokine production between vaccine groups also did not clearly identify any cytokines that correlated with protection (see Supplementary Table S1). For example, higher levels of TNF-α were present in the spleens of unvaccinated mice than BCG vaccinated mice (Fig 4C) and even though BCG vaccination resulted in significantly more IFN-γ in draining lymph nodes compared to unvaccinated mice, this was not observed for ER + R_4_Pam_2_Cys vaccinated mice (Fig 4D).

We therefore based our analysis on comparisons between diseased or protected mice irrespective of the vaccines they received (Fig. 4E,F). Herein, we identified significant increases of IL-2, IL-6, IL-10 IL-17A, IFN-γ, MIP-1b, TNF-α in the lymph nodes of protected mice (Fig. 4F) and significant increases of IFN-γ, IL-6, and TNF-α in the spleens of diseased mice (Fig. 4E). While these data implicate these cytokines as correlates of protection and disease, it did not rank the importance of each cytokine towards either outcome.

### Identifying immune responses that predict vaccination outcome

To identify the immune parameters (features) that associate with the response variable ‘vaccine protection’ (here measured as time to reach our bioluminescence detection threshold) independent of the vaccine used, we conducted a univariate regression analysis using the Cox proportional hazards model. We used this model as it accounts for the fact that a subset of mice (observations) was right-censored, as vaccination outcomes were not measured after 24 weeks post challenge. For each of the 28 immunological features, their association with the response variable (time-to-bioluminescence measured in weeks) was ranked using concordance index (CI) scores. The CI is analogous to the area under the ROC curve, with a CI value of 0.5 indicating a random correlation and 1 indicating a perfect, positive correlation (62). The CI for each univariate regression analysis was used to rank the strength of association for each of the 28 features against the response variable. Using a CI cut-off of 0.70, the top six features were identified as well as the direction of their association with prevention of development of bioluminescence (Table 2). Low levels of IFN-γ and high levels of IL-2 produced in mouse splenocytes were the top two immune parameter features influencing this model, reflected in the individual correlation between their respective titres and time-to-bioluminescence (Fig. 5A, B).

**Table 2:** High-scoring immune features associated with delayed bioluminescence.

| Feature number | Feature | Tissue site | Association with delayed bioluminescence | Concordance index |
| --- | --- | --- | --- | --- |
| 1. | IFN-γ | spleen | negative | 0.786 |
| 2. | IL-2 | spleen | positive | 0.769 |
| 3. | IL-6 | spleen | negative | 0.759 |
| 4. | TNF-α | spleen | negative | 0.745 |
| 5. | IL-2 | lymph node | positive | 0.715 |
| 6. | IFN-γ CD4<sup>+</sup> T-cell | spleen | negative | 0.707 |

**Figure 5.**
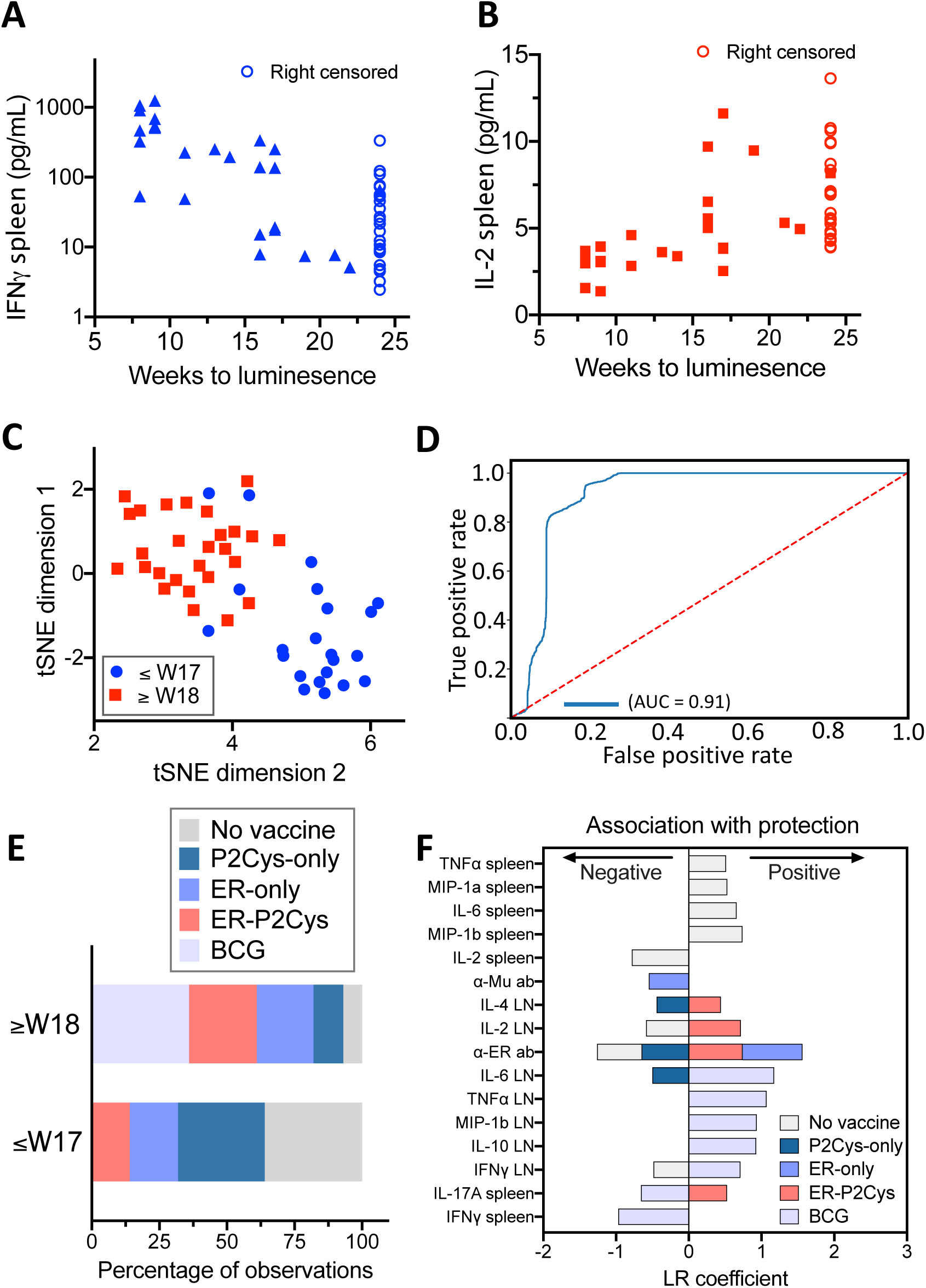
Statistical modelling to identify immune parameters (features) associated with vaccine protection. Univariate Cox Proportional hazards models were specified for each of the 28 immunological features to test their association with the response variable (time-to-bioluminescence measured in weeks). The resulting concordance index (CI) scores were obtained and the six features with a CI >0.7 were retained. The inverse associations of the top two features **(A)** IFN-γ and **(B)** IL-2 produced in murine splenocytes at the experimental end-point. **(C)** Plot depicting a two-dimensional representation of the top six features that associate with time-to-bioluminescence from the unsupervised t-SNE. The shapes/colours indicate the two groups identified through K-means clustering, of bioluminescence by 8-17 weeks or at 18 weeks and beyond (up to 24 weeks). **(D)** Receiver operator curve (and corresponding area under curve), displaying the trade-off between sensitivity and specificity across all thresholds for 1,000 random train-test splits of a logistic regression classifier (90% of observations used for training). The red dotted line depicts the expectation of a random classifier and the blue line depicts the model performance. **(E)** Proportion of observations across treatment groups for each of the classes both 8-17 weeks and 18-24 weeks and those with no detection. **(F)** Group-specific univariate logistic regression analyses for each of the five treatment groups. Model coefficients were used to determine both the strength and direction of association of each feature with that of each treatment group. Depicted are those features with a corresponding p-value<0.05 (Table S2).

Next, using these top six features, we performed unsupervised machine learning to reveal any structure without the influence of labels, such as arbitrarily imposed bioluminescence thresholds. Here, the data separated into two main clusters. K-means clustering was then applied to assign mice (observations) to the two cluster groups. Inspection of the resulting cluster membership with respect to time to bioluminescence showed a separation of the mice either side of 17 weeks post *M. ulcerans* challenge (Fig. 5C).

Given that the clusters identified through unsupervised learning closely resembled a temporal breakpoint at 17 weeks, we further investigated this binary divide in the data. We used multivariate logistic regression and developed a low-error classifier that could generalize to unseen data, using the underlying structure apparent in the immunological data. We then tested the classifier through extensive cross-validation. Observations (mice) with bioluminescence above threshold between weeks 8-17 were assigned a ‘0’ and ‘1’ for those with bioluminescence detected in weeks 18-24 or no detection throughout the experiment period. The model included the top six features (Table 2) and was validated through 1,000 random train test splits, with 90% of observations comprising the training groups at each split. The resulting classifier probabilities were used to calculate the area under the ROC curve, AUC = 0.91 (True negatives = 1774, True positives = 2662, False negatives = 120, False positives = 444). The low error and generalizable nature of this classifier demonstrates the existence of a robust structure in the data, in the form of two clusters separated around week 17 (Fig. 5D) and highlights the strong association of the six identified immune parameters with outcome. Most notably here it appears that tissue specific immune responses are important, with a correlation between the appearance of a tail ulcer and evidence of a systemic (spleen) responses and protection correlating with both local (draining lymph nodes) and systemic (spleen) response (Table 2). This association of BU and the production of inflammatory cytokines (IL-6, IFN-γ and TNF) as possible markers of infection, indicated that we could also identify correlates of immune protection against BU.

### Assessment of vaccine-specific immune responses

To dissect the immune responses associated with ER + R_4_Pam_2_Cys vaccination versus BCG and the different controls we noted the differences in the percentage of observations (mice) belonging to specific treatment groups between the two clusters observed above (Fig. 5C) separated by week 17. This summary was reflective of the survival analysis and showed that unvaccinated mice and those that received the adjuvant or ER alone were predominantly in the weeks 8-17 cluster, while ER + R_4_Pam_2_Cys and BCG-vaccinated mice were predominantly in the weeks 18-24 cluster (Fig. 5E). In order to obtain the individual immune profile of the different treatment groups, group-specific univariate logistic regression analyses were undertaken. Here, the response variable was coded as a ‘1’ for membership in a particular group and ‘0’ for membership in all other groups. Analyses were conducted for each of the five treatment groups and the model coefficient weights were used to determine both the strength and direction of association of each feature with that of each treatment group. The resulting *p*-values from the model coefficients were used to assess significance of the associations (Fig. 5F, Supplementary Table S2). The combination of the ER antigen and R_4_Pam_2_Cys together were important for inducing protective responses that associated with local production of IL-4, IL-2, IL-17A in the DLN, in addition to ER-specific antibody responses. Neither ER antigen or R_4_Pam_2_Cys alone induced this profile and protection using the former was only linked to ER-but not *M. ulcerans*-specific antibody responses (post-challenge timepoints). In comparison, BCG vaccination-mediated protection was associated with a greater breadth of localised cytokines responses than ER + R_4_Pam_2_Cys, with higher IL-6, TNF-α and MIP-1β in the DLN. Of note, evidence of systemic inflammation, such as IL-17A and IFN-γ in the spleen was associated with poorer BCG performance.

## Discussion

In this study, we investigated various immune parameters induced by the use of BCG and an experimental subunit vaccine against BU and sought to identify immune correlates associated with protection in a mouse tail infection model. Studies have shown that the tail is a suitable location for infection as BU predominantly affects extremities (11, 52, 63) and in combination with the use of bioluminescent *M. ulcerans*, offers several advantages over footpad, hock or ear infections used in most BU vaccine studies (as shown in Table 1 and (64)). Tail infections are less likely to affect mouse mobility, or cause rapid tissue loss and may also prevent added trauma, inflammation or secondary infections at the challenge site (65). A key feature of this model also allows the use of a significantly lower bacterial challenge dose compared to other studies (approximately 14-20 CFU compared to 10^4^-10^6^ CFU) (25, 32-37, 49, 52-54) (Table 1) and enables measuring bacterial growth in the same animal over time. This lower dose is likely to be more physiologically relevant, in terms of reflecting the bacterial inoculum that occurs during *M. ulcerans* transmission to humans (10, 11, 59). Sporadic healing of BU disease was also seen in this model, an observation that has been noted in humans and other animals (29, 66-68).

Vaccination with the ER + R_4_Pam_2_Cys formulation resulted in protection of 60% of mice from BU challenge. Although more of these vaccinated mice reached our defined disease outcome after 24 weeks compared to BCG-vaccinated mice, the difference was not statistically significant. There was also no significant difference in the number of mice displaying any bioluminescence between each group. The fact that ER + R_4_Pam_2_Cys was significantly more protective than vaccination with R_4_Pam_2_Cys alone indicates any non-antigen specific triggering of innate immune responses by this adjuvant (69, 70) (71) was not sufficient for conferring long-lasting protection and the inclusion of the ER protein was necessary to achieve any protective effects. This was further evident by vaccinating with ER alone. Despite not seeing any clear significant differences in the clinical outcomes at the end of the trial when compared to ER + R_4_Pam_2_Cys vaccination, the inclusion of R_4_Pam_2_Cys delayed disease onset by ∼8 weeks and correlated with the induction of significantly more ER-specific antibodies after a primary and booster vaccination.

The presence of these antibodies was nonetheless insufficient to provide total protection against *M. ulcerans* challenge. This is perhaps not unexpected given that other studies have also reported little correlation between strong BU antibodies and protection, in mice (34) or sera of BU patients (57, 72). To identify other immune correlates associated with the protective effects observed in our study, a Cox proportional hazards regression model was utilized. Here, the univariate analyses allowed us to identify the top six features that most strongly associated with differences in time to detection of bioluminescence. These models assume normalized data, and although we cannot infer how much an increase or decrease in units could affect the clinical outcome, we are able to rank each factor based on its contribution to either disease or protection. This model also considers the effect of each factor in delaying the onset of disease rather than just modelling through a binary ‘protected’ or ‘diseased’ outcome. Similar regression modelling has been described to predict outcomes for tuberculosis patients after treatment, effect of hospital-acquired *Clostridium difficile* on hospital stay and survival of *Staphylococcus aureus* in milk (73-75). The model assumes that eventually all mice will succumb to disease and due to the constraints in our data, it cannot determine threshold levels of cytokines that will predict disease outcomes.

The cytokines most associated with protection in BCG mice were different to those identified in mice vaccinated with ER+R_4_Pam_2_Cys, which is not surprising given that BCG is a multi-antigen live-attenuated vaccine and thus likely to utilise both common and distinct protective responses and mechanisms to those induced by our vaccine candidate.

Through multivariate logistic regression modelling we identified the presence of IL-2 in the spleen and lymph nodes as markers that were most strongly associated with protection. Although there are no studies that directly link IL-2 with protection against *M. ulcerans*, it plays a key role in the differentiation, proliferation and maintenance of T cell responses (76). Therefore, it is perhaps not surprising that its role in this murine model is likely to be important for the induction of protective adaptive immunity against *M. ulcerans.*

However, our results did not show any correlation between levels of cytokine producing CD4^+^ and CD8^+^ T cells and protection in vaccinated groups despite studies that have shown them to play a role in BU control (77, 78). In particular, *M. ulcerans-*specific CD4^+^ T cells have been found to migrate to the site of infection from draining lymph nodes early in infection but are depleted as the infection persists (77), an effect that could be attributed to the ability of the *M. ulcerans* exotoxin mycolactone to impair T cell and macrophage function (41, 44, 78).

In addition, we also identified the presence of systemic IL-6, TNF-α and IFN-γ (in the spleen) to be strongly associated with disease. IL-6 is a pro-inflammatory cytokine produced by many cell types in response to pathogens and is linked to the production of TNF-α, both of which can be detected in BU lesions and serum of BU patients (78, 79) (80). TNF-α in particular, plays a key role in inflammatory cell recruitment and in conjunction with IFN-*γ*, increases the phagocytic ability of macrophages to enhance killing of mycobacteria (81, 82). However many studies have shown that mycolactone suppresses TNF-α production by T cells and especially macrophages (83, 84), decreasing their ability to control BU infection (85).

IFN-*γ* itself has also been shown to be important for controlling *M. ulcerans* infection as IFN-*γ* deficient mice cannot prevent the onset of disease (86). It is also detected at high levels in patients with both developed ulcers and early lesions (87) and healed ulcers (88) where it is believed to mediate macrophage function (89) and drive iNOS expression to facilitate bacterial killing (90). Altogether, the fact that these cytokines are elevated at a systemic level in the diseased animals in our study but not in lymph nodes draining from the tail suggests that their activity is being dampened at the site of infection. These effects do not appear to be present in protected animals where increased levels of cytokines are detected in the draining lymph node and not the spleen. In fact, the localised, but not systemic presence, of these and other cytokines including IL17A, MIP1b and IL10 are strongly associated with protection.

Although most of the immunosuppression during BU infection can be attributed to mycolactone, chronic inflammation can also be key driver as noted by the increased splenic cytokine levels. Many cell types have been implicated as the cause of immunosuppression in cancers, chronic viral infections (such as HCV, HIV, HBV) and even *M. tuberculosis* infection. These include myeloid-derived suppressor cells (MDSCs) (91) regulatory T cells (T_reg_) (92) and T helper 17 (T_h_17) (93), which can suppress effector T cell function and inhibit NK and dendritic cell activity through direct cell-to-cell interactions or the production of immunosuppressive cytokines. MDSCs in particular can be recruited by IFN-γ (94) and IL-6 (95), both of which are found in higher levels in our unprotected mice. On the other hand, while T_h_17 cells are crucial for the control of infection, especially extracellular bacterial and fungal infections, elevated frequencies can lead to tissue inflammation alongside matrix destruction, autoimmunity and vascular activation (96). The observation that higher systemic IL-17A correlates with the lack of protection suggests that these cells play a role in determining BU disease outcomes.

Tissue changes due to chronic infection could also play a compounding effect on the severity of BU disease outcomes. Our histological analysis of BU-infected tail tissue showed a loss of muscle and epidermis, changes in connective tissue and loss of vasculature which may explain why lymphocytes and other immune cells are unable to access the sites of greatest infection and tissue damage. BU tissue necrosis can also extend some distance from the site of bacterial colonisation, an observation that led to the identification of mycolactone as the cause of coagulative necrosis (39, 97, 98). Mycolactone has been well described as causing cell death to skin-resident cells such as fibroblasts, adipocytes, keratinocytes and endothelial cells (39, 99, 100). Primary human dermal microvascular endothelial cells are especially sensitive to mycolactone and after exposure lose their ability to activate a key anticoagulant protein (protein C) after exposure, causing a reduction in intravascular fluidity and preventing immune cell infiltration to the infection site (99). Thus, the combination of immunosuppressive immune host responses and tissue destruction, in conjunction with mycolactone at the site of infection, may increase the risk for poorer disease outcomes for those chronically infected with BU.

Although we have identified several factors associated with disease and protection, our results provide impetus to further expand these profiles and establish their importance. For example, changes in cytokines levels before challenge and throughout the infection phase could be monitored and integrated into models, as well as analysing frequencies of various other innate- and adaptive-immune cell populations and identifying those that produce cytokines of interest. In evaluating and demonstrating that a subunit vaccine can protect against BU in our mouse challenge model, albeit not as efficacious as BCG, our results showed that protection can be mediated through different immune mechanisms. Disease progression was also commonly linked to the presence of pro-inflammatory cytokines in the spleen and not the lymph node. These profiles indicate that localised and not systemic responses are more important for conferring protection and also provide a template that could guide the design and development of novel vaccination strategies against BU.

Finally, we conclude that the mycolactone biosynthesis pathway constitutes a viable vaccine target to protect against *M. ulcerans*. As *M. ulcerans* is slow-growing and requires its highly conserved mycolactone PKS for virulence, the development of resistance is unlikely. As such, approaches based on the use of multiple PKS enzymatic domains may prove even more efficacious. Moreover, studies that have introduced *M. ulcerans* and *M. marinum*-specific proteins into BCG have been shown to increase its protective effect. Collectively, this demonstrates the additive power of using a broader suite of antigens and the potential for a viable vaccine against BU.

## Methods

### Strains and culture conditions

*Escherichia coli* ClearColi©BL21 (DE3) containing the plasmid, pJexpress-ER (strain TPS847) was grown at 37°C in Luria-Bertani (LB) broth (Difco, Becton Dickinson, MD, USA) supplemented with 100 µg/ml ampicillin (Sigma-Aldrich, USA) to express the enoyl reductase (ER) protein (57). Log-phase bioluminescent *Mycobacterium ulcerans* (strain JKD8049 containing integrated plasmid pMV306 *hsp:lux*G13) (10, 60) was grown at 30°C in 7H9 broth or 7H10 agar (Middlebrook, Becton Dickinson, MD, USA) supplemented with oleic acid, albumin, dextrose and catalase growth supplement (OADC) (Middlebrook, Becton Dickinson, MD, USA), 0.5% glycerol (v/v) and 25 µg/ml kanamycin (Sigma-Aldrich, USA). *M. bovis* BCG (strain ‘Danish 1331’) used for vaccinations was grown at 37°C in 7H9 broth or 7H10 agar supplemented with OADC. Mycobacterial colony counts from cultures or tissue specimens were performed using spot plating as previously described (10). All culture extracts were screened by LC-MS for the presence of mycolactones as previously described to ensure bacteria used in transmission experiments remained fully virulent (101).

### Recombinant protein expression

Overnight culture of *E. coli* TPS847 was diluted to OD_600_ = 0.05 in LB broth. The culture was incubated at 37°C with shaking at 200 rpm until OD_600_ = 0.6-0.7, followed by the addition of 1 mM IPTG (Isopropyl b-D-1-thiogalactopyr-anoside) to induce protein expression for a further four hours. Cells were then resuspended in wash buffer (8 M urea, 150 mM sodium chloride, 10% glycerol) and sonicated at amplitude 60 (QSonica Ultrasonic Liquid Processor S-4000, Misonix) until the solution turned clear. The lysate was filtered with a 0.22 µM filter (Millipore) to remove cellular debris and protein was column-purified using anti-histidine resin (ClonTech). The resin was washed with wash buffer which was gradually replaced with tris buffer (20 mM Tris-HCl, 150 mM sodium chloride, 10% glycerol) over ten washes followed by two washes with tris buffer containing 20 mM imidazole. Protein was eluted in tris buffer containing 200 mM imidazole and dialysed in phosphate buffered saline (PBS) before concentration using a microcon column (Millipore). Proteins were tested for endotoxin contamination using Pierce™ limulus amoebocyte lysate assay (Thermo Scientific™) and relative size was confirmed by sodium dodecyl sulphate polyacrylamide gel electrophoresis.

### Sodium dodecyl sulphate polyacrylamide gel electrophoresis (SDS-PAGE)

Samples were denatured in an equal volume of 2 x sample loading buffer (40% (v/v) 0.5M Tris-HCL pH6.8, 10% glycerol, 1.7% (w/v) SDS, 10% 2-B-mercaptoethanol, 0.13% (w/v) bromophenol blue in distilled water) at 100°C for 5 minutes. Ten microlitres of each sample and SeeBlue® Plus2 pre-stained protein standard (Invitrogen) was loaded onto a 0.5mm 12% polyacrylamide gel under reducing conditions, as previously described (102). The gel was run in buffer containing 0.3% (w/v) Tris, 1.44% (w/v) glycine and 0.1% (w/v) SDS in distilled water for 1 hour at 150 volts (Mini-protean vertical electrophoresis cell, Bio-Rad), stained in Coomassie stain (45% methanol, 10% acetic acid 0.25% (w/v) Coomassie brilliant blue in distilled water) for 1 hour and destained in Coomassie destain (33% Methanol, 10% acetic acid, 60% distilled water) until protein bands were visualised.

### Western Blotting

Protein separated on a 12% polyacrylamide gel was transferred to a nitrocellulose membrane in a tris-glycine transfer buffer (1.5 mM Tris, 12mM glycine, 15 % methanol (v/v) in distilled water) for 1 hour at 100 volts (Mini Trans-Blot Cell, Bio-Rad) and incubated in blocking buffer (5% (w/v) skim milk powder and 0.1% Tween-20 in PBS) overnight at 4°C. The membrane was then incubated with anti6xHIS-HRP antibody (Roche Applied Science, USA) at 1:500 dilution for 2 hours and washed in PBS containing 0.1% Tween-20 prior to exposure to developing solution (Western Lighting Chemiluminescence kit, Perkin Elmer, USA) according to the manufacturer’s guidelines. Chemiluminescence was detected using an MF ChemiBIS gel imaging system (DNR Bio-Imaging Systems, Israel).

### Particle size analysis of protein antigen and lipopeptide formulations by dynamic light scattering (DLS)

The association between protein and R_4_Pam_2_Cys was measured using dynamic light scattering (DLS) by mixing 5 µg of protein with increasing amounts of lipopeptide in 50 µl PBS. The size distribution of particles in solution (presented as hydrodynamic radius) were measured in 4µl of cyclin olefin co-polymer cuvettes using a DynaPro NanoStar DLS instrument (Wyatt Technology, CA, USA) equipped with 658nm laser with a scattering angle of 90°. Measurements were acquired in triplicate with each measurement consisting of 30 readings at 5 second intervals at 25°C. Data was analysed using Dynamics software (v7.1.7.16).

### Vaccination of animals

The synthesis and purification of the branched cationic lipopeptide, R_4_Pam_2_Cys, was performed as previously described (103, 104). Each vaccine dose contained 25 µg protein formulated in PBS with R_4_Pam_2_Cys at a 1:5 molar ratio of protein to lipopeptide in a final volume of 100 µl. Live-attenuated *M. bovis* BCG strain ‘Danish 1331’ was grown to log phase and stored at −80°C in 20% glycerol until used. Bacteria were washed with PBS and resuspended in 200ul, before administration at 4.7 × 10^5^ bacteria per dose. All vaccines and control formulations were sonicated for 5 minutes in a waterbath sonicator before being administered.

Female 6-week old BALB/c mice were sourced from ARC (Canning Vale, Australia) and housed in individual ventilated cages. Food and water were given *ad libitum*. Experiments were approved by The University of Melbourne Animal Ethics Committee (Approval identification number: 1613870). For vaccination using R_4_Pam_2_Cys, animals were inoculated subcutaneously at the base of tail (100µl per dose at 50 µl per flank) and boosted 21 days later with the same formulations. Mice vaccinated with *M. bovis* BCG were given one dose subcutaneously in a similar manner (200 µl per dose at 100µl per flank). There were 10 mice in each vaccination group.

### M. ulcerans challenge

Mice were challenged with bioluminescent *M. ulcerans* on day 35 as described previously (10). Briefly, tails of isoflurane anaesthetised mice were dipped in 7H9 culture containing log-phase bioluminescent *M. ulcerans* bacteria (concentration 1.27 × 10^6^ CFU/mL (range: 1.07×10^6^ – 1.46 ×10^6^ CFU). Contaminated tails were then pierced once subcutaneously with a sterile 25-G needle. The infectious dose was calculated to be 17 CFU (range: 14-20) using methods previously described (10). Mice were allowed to recover and monitored for up to 24 weeks after infection and sacrificed when tail ulceration was observed wherein spleens, lymph nodes and sera were harvested for immunological analysis.

### IVIS imaging

Infected mice were imaged weekly from 6-weeks post-infection to detect the emission of bioluminescence. Images were captured using the Lumina XRS Series III In Vitro Imaging System (IVIS®) (Perkin Elmer, MA, USA) and Living Image Software v3.2 with the following settings: Field of View 24, relative aperture f’1.2, medium binning, 60s exposure. Bioluminescence was calculated using Living Image Software v3.2.

### Serum antibody titre measurements

Serum antibody titres were measured by enzyme linked immunosorbent assay (ELISA) (105) using plates (Nunc, Thermo Scientific) that were previously coated with antigen overnight, either purified recombinant ER protein or heat-killed whole cell *M. ulcerans* lysate. The presence of bound antibodies were detected by incubating serum-exposed wells with horse radish peroxidase conjugated rabbit anti-mouse IgG (Dako, Glostrup, Denmark) for 2 hours followed by the addition of the enzyme substrate (0.2mM ABTS in 50mM citric acid containing 0.004% hydrogen peroxide and left to develop for 10-15 minutes before the addition of 50nM sodium fluoride to stop the reaction. Plates were read at dual wavelengths of 505 and 595 nm on plate reader (LabSystems Multiskan Multisoft microplate reader) and antibody titres expressed as the reciprocal of the highest dilution of serum required to achieve an optical density of 0.2.

### Intracellular cytokine staining

Single cell suspensions were derived from the spleen and draining lymph nodes and resuspended in RP10 media (RPMI 1640 (Sigma) supplemented with 10% foetal bovine serum (Gibco, ThermoFisher Scientific, Waltham, MA USA), 2mM L-glutamine, 1mM sodium pyruvate, 55 µM 2-mercaptoethanol, 12 µg gentamycin, 100 U/ml penicillin and 100 µg/ml streptomycin). Spleen and lymph node-derived cells were cultured in 96-well plates (CoStar, Corning, USA) at 1 ×10^7^ cells/per well and 1 ×10^5^ cells/well, respectively 200 µl of RP10 containing 10 U/ml IL-2 (Roche, Mannheim, Germany), 1µg/ml plate-bound anti-CD28 (BD Pharmingen, Becton Dickinson, Clone 37.51) and 20 µg/ml ER protein for 12 hours at 37°C in 5% CO_2_. Golgiplug (1µg/ml) (Becton Dickinson) was added for the last 4 hours of incubation. Cells were then stained with 7AAD-Live/Dead stain dye (Biolegend, CA, USA), BV510-anti-B220 (BD Horizon, Becton Dickinson, Clone RA3-6B2), BV605-antiCD4 (Biolegend, Clone RM4-5), APC-Cy7-anti-TCRb (BD Pharmingen, Clone H57-597) and PE-Cy7-anti-CD8 (BD Pharmingen, Clone 53-6.7) anti-mouse monoclonal antibodies at 4°C in the dark. Intracellular staining was performed by fixing cells with Cytofix/Cytoperm solution (Becton Dickinson, USA) followed by permeabilisation and intracellular staining with Perm/Wash buffer (Becton Dickinson) and BV786-IFN-γ (BD Horizon, Clone XM G1.2), AF647-IL-17A (BD Pharmingen, Becton Dickinson, Clone TCII-18H10) and PE-TNF-α (BD Biosciences, Clone MP6-XT22) antibodies for 30 minutes at 4°C before analysis on an LSR Fortessa flow cytometer (BD Biosciences, US). Data analyses were performed using FlowJo (Tree Star, OR, USA).

### Cytokine Bead Array

Spleen and lymph node-derived cells were incubated in 500 µl RP10 supplemented with 25 µg/ml ER protein for 72 hours at 37°C in 5% CO_2_. Supernatant was collected and a cytokine bead array was performed using a mouse flex set (BD Biosciences, USA) to detect IL-2, IL-4, IL-6, IL-10, IL-12/IL-23p40, IL-17, IFN-γ, TNF, MCP-1, MIP1α, MIP1β as per the manufacturer’s instructions. Samples were acquired using a FACSCanto II flow cytometer (BD Biosciences) and cytokine quantities calculated using FCAP Array™ Software v3.0.

### Histology and microscopy

Tail tissues from the site of infection were fixed in PBS containing 10% non-buffered formalin then embedded in paraffin and sliced into 10 µM thick segments. The sliced segments were Ziehl Neelsen- or H&E-stained prior to microscope imaging. Images of tail segments were captured using a light microscope (Olympus BX53 Light microscope, Olympus-Life Science).

### Statistical analysis

GraphPad Prism software (GraphPad Software v7, CA, USA) was used to perform statistical analyses on the antibody titre, time to luminescence, T cell numbers and cytokine titre data. Antibody titres were analysed using one-way ANOVA with Tukey’s correction for multiple comparisons. The time to bioluminescence data was displayed as a Kaplan-Meier plot and differences determined using a Log-Rank (Mantel-Cox) test. Mann-Whitney tests were performed to compare cytokine titres between protected and diseased mice and for comparisons between vaccination groups. All tests were conducted at the 5% significance level.

### Statistical modelling

Twenty-eight data features (*i.e.* the immune parameters measured in each mouse, refer to Table S1) were transformed using the R package *bestNormalize* (106). Transformed features were then normalised (between 0 and 1 for each feature) using the *MinMaxScaler* function of *Scikit-learn* (107). As many of the vaccination outcome observations (time-to-bioluminescence) were right-censored, we employed the Cox proportional hazards regression analysis using the *Scikit-survival* module of *Scikit-learn* in Python (108). Here, univariate analyses were run for each of the 28 features using the continuous response variable of time-to-bioluminescence. The standard metric for assessing the predictive performance of a survival model is the concordance index (CI) (62, 109, 110). A CI >0.7 was used to identify the top six features of this model. Unsupervised learning and data dimensionality reduction is an ideal way to identify structure in continuous data without the influence of labels. The method of *t-Distributed Stochastic Neighbor Embedding* (t-SNE) is a dimensionality reduction technique that retains both the global structure and local layout of the high-dimensional data through exchanging the Euclidean distances between all pair of data points into heavy-tailed conditional probabilities (111). This method is advantageous over conventional principal component analysis (PCA), as it does not rely on a linear assumption and can capture nonlinear relationships (111). We explored the data, independent of labels, by reducing the top six features obtained from the Cox proportional hazards regression analysis to a two-dimensional space using the t-SNE package in *Scikit-learn* (107). The two clusters detected through visual inspection were objectively defined, with observations assigned to two groups using K-means clustering, as implemented in *Scikit-learn* (107). A multivariate logistic regression classifier was then built using the top six features, with the two clusters identified by *t-SNE* as the response variables. To reduce the possibility of over-fitting, the model was validated through 1,000 random train-test splits, in which 90% of the observations made up each training set. These models were built using the logistic regression classifier as implemented in *Scikit-learn* (107) and Receiver-Operator-Characteristic curves were used to evaluate model performance (112). In order to assess the immune features that were associated with different vaccination groups, a univariate logistic regression analysis was then conducted for each group using *R* (112). The estimated model coefficients were used to assess the direction and strength of the association, and the corresponding p-value used to determine statistical significance at the 5% significance level.

## Acknowledgments

We thank Laura Leone for expert assistance with histology. This research was supported by the National Health and Medical Research Council, Australia (GNT1008549). The funders had no role in study design, data collection and interpretation, or the decision to submit the work for publication.

## Supplementary data

**Table S1** - Vaccination data

**Table S2** - Summary of group-specific univariate logistic regression coefficients

